# Tree House Explorer: A Novel Genome Browser for Phylogenomics

**DOI:** 10.1101/2022.01.03.474706

**Authors:** Andrew J. Harris, Nicole M. Foley, Tiffani L. Williams, William J. Murphy

## Abstract

Tree House Explorer (THEx) is a genome browser that integrates phylogenomic data and genomic annotations into a single interactive platform for combined analysis. THEx allows users to visualize genome-wide variation in evolutionary histories and genetic divergence on a chromosome-by-chromosome basis, with continuous sliding window comparisons to gene annotations, recombination rates, and other user-specified, highly customizable feature annotations. THEx provides a new platform for interactive phylogenomic data visualization to analyze and interpret the diverse evolutionary histories woven throughout genomes. Hosted on Conda, THEx integrates seamlessly into new or pre-existing workflows.

## Introduction

In recent years, rapid advancements in massively parallel sequencing technologies have enabled researchers to collect large volumes of genome-scale phylogenetic and genomic data for many species. Computer software designed to analyze these data have been available to users for more than a decade, but the separation of phylogenetic and genomic visualization tools has remained a common challenge for evolutionary biologists. IGV and the UCSC Genome Browser are important tools for visualizing genomic data, while Iroki, FigTree, and ETE (v3) are used to visualize and manipulate large phylogenetic trees separately (Kent et al. 2002; Rambaut 2006; Thorvaldsdottir et al. 2013; Huerta-Cepas et al. 2016; Moore et al. 2020). These programs have become indispensable for interrogating specific features of genomic and phylogenetic data separately, but there has yet to be a movement toward combined analysis until recently.

Genomes are mosaics of evolutionary histories that reflect ancient signatures of species divergence, as well as incomplete lineage sorting (ILS) and gene flow. Understanding how and why phylogenetic signal varies across species genomes can yield powerful insights into evolutionary histories and adaptive evolution. By integrating diverse data types with local genealogies, one can more readily differentiate genetic variation that is consistent with the species tree from that stemming from natural selection, ILS, or gene flow (Figueiro et al. 2017; Edelman et al. 2019; Li et al. 2019; Small et al. 2020; Hennelly et al. 2021; Nelson et al. 2021). However, a tool has yet to be developed that can simultaneously analyze phylogenetic signal variation with other chromosomal and gene-based annotations (i.e. a phylogenomic browser).

Here we present Tree House Explorer (THEx), a novel genome browser designed to explore phylogenetic signals in parallel with chromosomal and gene-based annotations in an all-in-one application. THEx offers two different dashboards, Tree Viewer and Signal Tracer, that provide highly interactive and customizable graphing experiences that simplify the exploration of diverse phylogenetic histories and window-based calculations in an annotated genomic context. Exploring data from local and genome-wide views makes the identification of complex divergence patterns easier to identify and offers a unique way to connect phylogenetic signal with underlying genomic data like recombination rate, gene annotations, divergence time estimates, or any other window-based data type. By facilitating synchronous visualization of phylogenetic signal and additional data types, THEx allows users to visualize more complex genomic associations that were previously hidden and place them in a genomic context. In addition to synchronous visualization, THEx enables users to download publication-ready figures in several different file types (e.g. .svg, .jpeg, .png).

THEx was developed using Python, Plotly’s open-source graphing library, and Dash, a web data analytical application framework. Deployed on Conda, an open-source package and environment management system, THEx is easily integrated into pre-existing pipelines and removes the complexities of gathering required software dependencies and ensuring correct version compatibility. THEx was tested using more than 200 mammal whole genome alignments and can comfortably analyze phylogenomic datasets containing hundreds of taxa on a local workstation (Genereux et al. 2020). Future development aims to improve its capabilities on large supercomputing clusters and servers. Documentation, example data sets, and tutorials can be found on the THEx GitHub page (https://github.com/harris-2374/THEx).

## New Approaches

### Overview

THEx comes with two dashboards, Tree Viewer and Signal Tracer, that provide unique approaches to investigating phylogenomic data. Tree Viewer is designed to investigate how phylogenetic signal varies across the genome with respect to various genomic data types (e.g., recombination, protein-coding regions, etc.). Signal Tracer is similar in that it provides a platform to investigate window-based calculations, but its focus is on displaying information per-taxon rather than per-topology. This distinction is vital because phylogenetic signal only shows general differences in the relationships among taxa, but does not provide specific information about what is different. Therefore, investigating information at a per-taxon level is required to understand the underlying reasons for changes in phylogenetic relationships. In addition to serving as a novel phylogenomics browser, THEx offers Linux and macOS users a custom companion toolkit called THExBuilder, which simplifies the generation and manipulation of THEx input files. Written in Python and deployed on Conda, THEx is straightforward to install and integrate into pre-existing data analysis pipelines. Overall, THEx takes the highly fragmented nature of phylogenomic analyses and simplifies the process by integrating phylogenetic signal and other chromosomal and gene-based data types (e.g., recombination rate, divergence time estimates, gene annotations, etc.) into a single browser for easy identification and interrogation of evolutionary distinct regions of the genome.

### THExBuilder Command Line Interface

THExBuilder is a growing toolset that helps users go from multiple-sequence alignments to THEx input files as quickly as possible while also providing tools that make working with and manipulating input files a straightforward task. Invoked by the command “*thexb*”, Linux and macOS users are provided a command-line suite of tools and pipelines that aid in generating and manipulating input files for THEx. THExBuilder offers one possible approach to generating a Tree Viewer input file from a multiple-sequence alignment Fasta file and provides several additional tools that enable users to manipulate Tree Viewer input files. For example, THExBuilder allows users to root or re-root all phylogenetic trees within a Tree Viewer input file to a new outgroup. This tool removes the need to alter the raw tree data and regenerate a new Tree Viewer input file. Another major improvement implemented within THExBuilder’s Tree Viewer pipeline is an improved phylogenetic tree binning algorithm called *topobinner*. Tree binning is a process in which a set of phylogenetic trees are grouped based on the relationships among taxa. Trees with identical relationships among taxa are grouped together and ordered high-to-low based on the number of trees within each group. This information is the basis of the visualization within Tree Viewer and is what allows users to explore the phylogenetic signal across their genomes. *Topobinner* is a user-friendly replacement for the legacy tree binning program, *PhyBin* (Newton and Newton 2013). It also provides a command to calculate raw divergence (p-distance) from a multiple-sequence alignment that can be viewed in Signal Tracer. THExBuilder comes a wide variety of tools and is under active development to improve and expand its capabilities.

### Tree Viewer Dashboard Interface

Tree Viewer is designed to integrate whole-genome phylogenetic signal with various chromosomal and gene-based data types. The dashboard is separated into two main sections; a navigation bar with collapsible menus (**Figure 1A & 1B**) and a multi-graph container (**Figure 1C & 1D**). The navigation bar contains five collapsible menus: an input/export menu, the main toolbar containing all data and graphing options, a summary-statistics menu, a graph customization menu, and a documentation section describing how to use the dashboard. Clicking Tree Viewer from the homepage will switch to the Tree Viewer dashboard and open a prompt for the user to select an input. Once the user selects the required input files and clicks submit, the data is checked for errors or incorrect formatting, and then the main toolbar is populated with the user-provided information.

**Figure 1.**
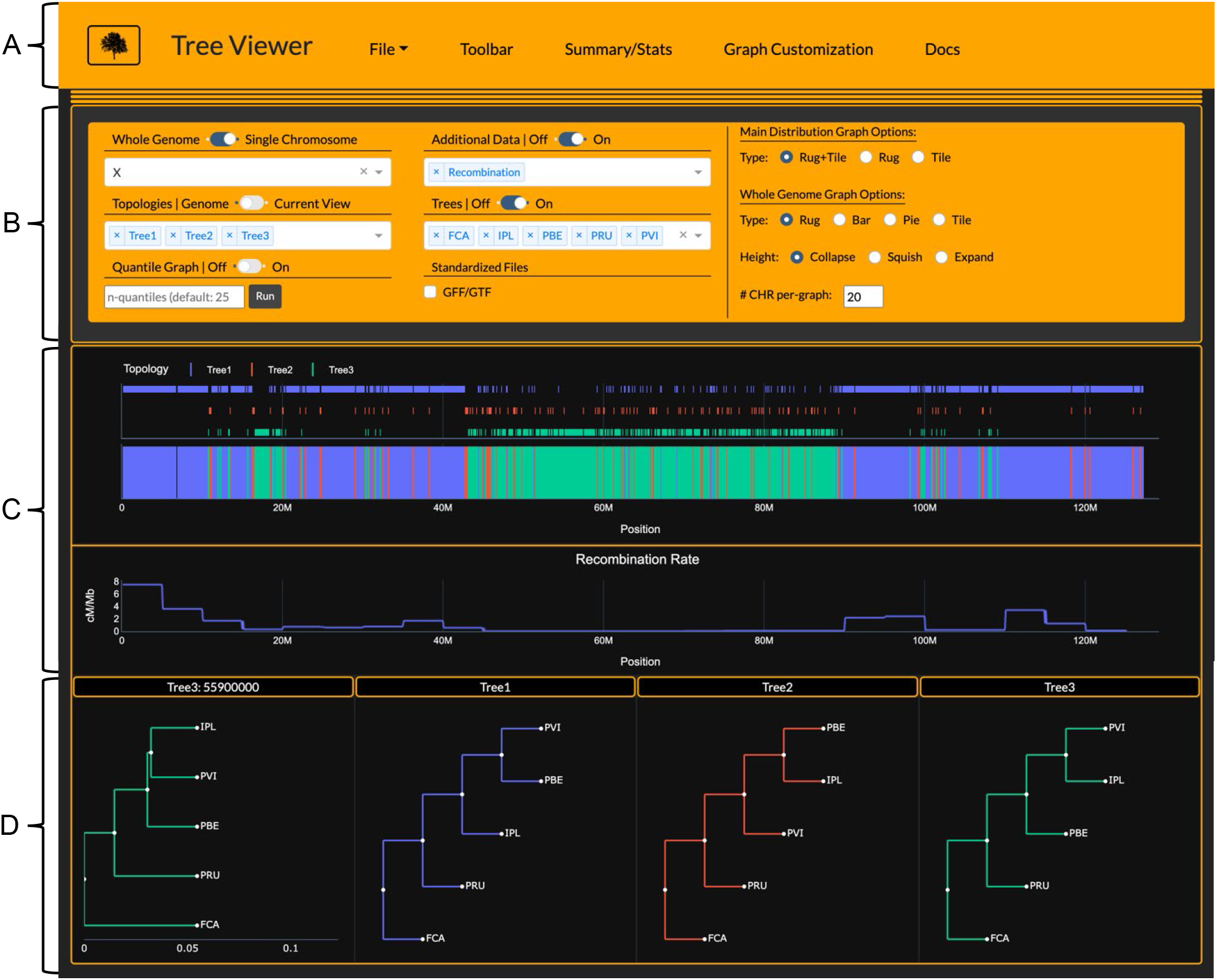
Screenshot of Tree Viewer interface displaying data of the Asian leopard cat lineage generated by Li et al. (2019). The navigation bar at the top of the page (A) contains all access points to the input/export options, main graph toolbar, summary/statistics, graph customization, and documentation. (B) shows the main toolbar where users control which graphs to display and what types of graphs to plot. (C) is the main graph container that shows Asian leopard cat lineage phylogenetic signal on top and recombination rate in the row below. (D) shows the user-selected tree topologies from the “Topologies” dropdown that are displayed across all loaded graphs. The first tree in (D) depicts a user-selected tree topology with respective branch lengths chosen by clicking on a window on the main phylogenetic signal distribution (top graph in C). The other three tree topologies are basic representations of each selected topology with unit branch lengths. Domestic cat (FCA), Rusty-spotted cat (PRU), Flat-headed cat (IPL), Fishing cat (PVI), and Asian leopard cat (PBE).

Once the data is loaded into Tree Viewer, users can explore their data either in whole-genome mode or single-chromosome mode. The viewing mode can be changed by toggling the switch between “Whole Genome” and “Chromosome” in the main toolbar (**Figure 1B**). Whole-genome view offers an overview of the selected topologies across the entire genome and can be graphed as a rug plot, bar plot, pie chart, or one-dimensional tile plot faceted by chromosome. Single-chromosome viewing mode enables users to zoom in and out of local regions of a chromosome to investigate discordance in direct comparison to additional data types selected in the “Additional Data” dropdown. The first graph to load shows the phylogenetic signal (variation in window-based trees or gene trees) across the chosen chromosome. If additional data types are also loaded, their x-axis range will sync to the main distribution graph (top graph of **Figure 1C**), allowing the user to zoom and pan across the genome while directly comparing multiple data types at once, such as recombination rate, divergence time estimates, or gene annotations.

Users can load basic representations of selected tree topologies with unit branch lengths and can optionally click on specific windows on the main distribution graph to view a specific window’s topology with branch lengths if they are present in the Newick trees within the Tree Viewer input file (**Figure 1D**). A dropdown menu is also provided under the “Trees” toggle, enabling users to prune trees down to a select number of taxa for easier visualization. This enables users to easily visualize sub-trees when the original input contains hundreds of taxa and does not alter the input file. Users can use the “File Pruning” export option in the “File” dropdown to prune trees down to a select set of taxa and re-bin topologies. This option allows users to select a subset of taxa and re-bin the resulting topologies either within THEx (<10 taxa and/or genome < 2.5Gb) or more efficiently, externally in the command line using THExBuilder’s *topobinner* script (>10 taxa and/or genome > 2.5Gb).

### Signal Tracer Dashboard Interface

Signal Tracer is a dashboard that allows for intuitive, chromosome and gene-aware investigation of variation in window-based calculations like genetic distance, branch length, or divergence times at single chromosome and whole genome views. Compared to Tree Viewer, Signal Tracer visualizes values per-taxa rather than per-topology. To demonstrate the functionality of Signal Tracer, we utilized previously published phylogenomic datasets from the cat family (Li et al. 2019). We generated separate Tree Viewer input files of the species from the Lynx and Asian leopard cat lineages and extracted per-taxa branch length information from the per-window Newick phylogenetic trees. By converting per-taxon branch length information into Signal Tracer’s tab-delimited input file format, we illustrate how users can visualize divergence between samples linearly across chromosomes rather than comparing hundreds-to-thousands of independent phylogenetic trees (**Figure 2**). This allows for simultaneous visualization and interpretation of underlying variation in the context of genic and chromosomal features and can highlight interesting deviations in genetic divergence, which otherwise may not be entirely obvious when looking at tree topologies alone.

**Figure 2.**
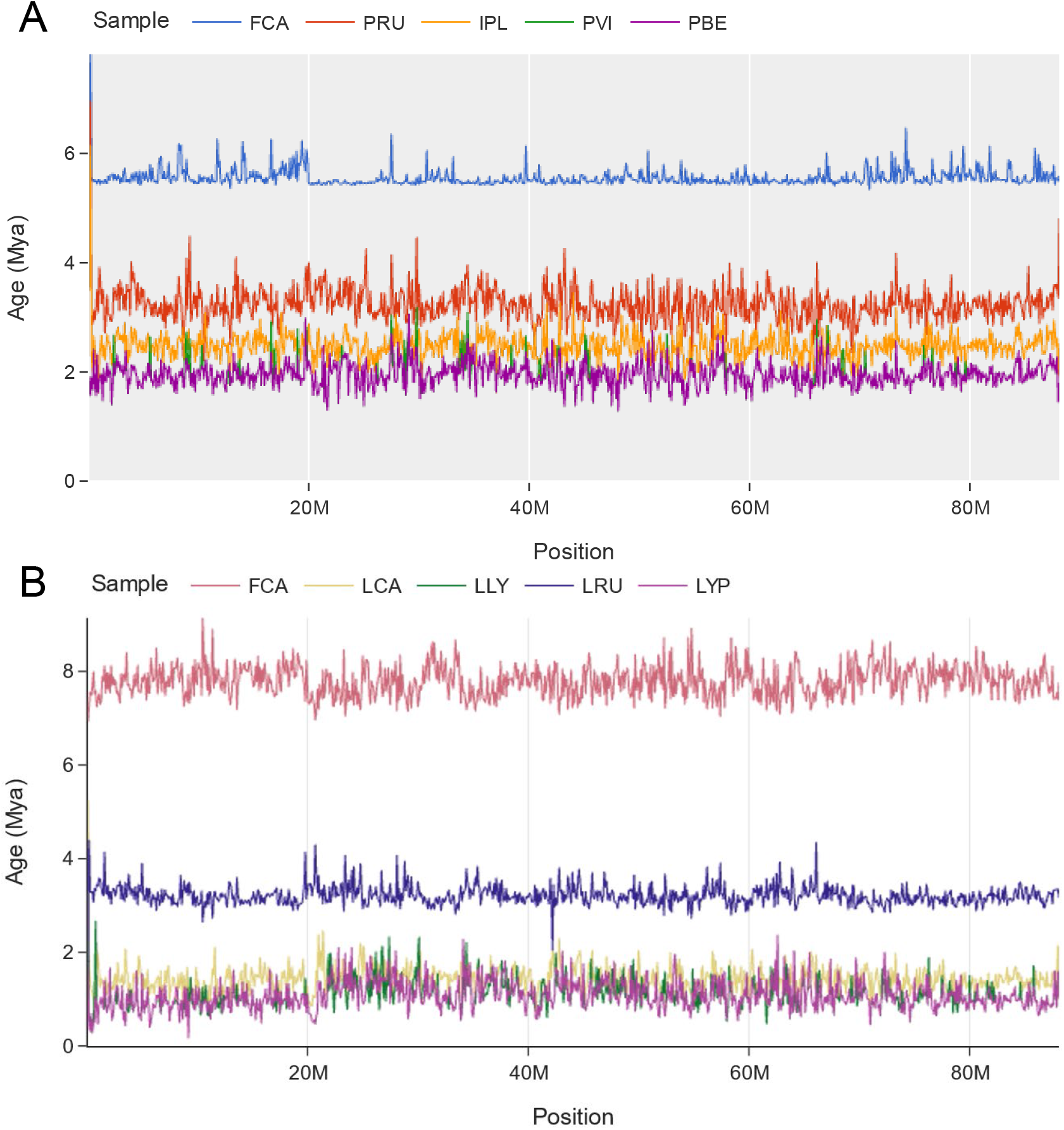
Signal Tracer plots of chromosome D2 for the Lynx (A) and Asian leopard cat (B) lineages with different themes and color palettes. Individual lines represent divergence time estimates (y-axis) of the Domestic cat (FCA), Canada lynx (LCA), Eurasian lynx (LLY), Iberian lynx (LYP), Bobcat (LRU), Rusty-spotted cat (PRU), Flat-headed cat (IPL), Fishing cat (PVI), and Asian leopard cat (PBE) across chromosome D2 (x-axis). Divergence time estimates for each taxon are derived from per-window phylogenetic trees in each lineage’s respective Tree Viewer input file.

For example, by comparing the branch lengths from phylogenies inferred for the Lynx (**Figure 2A**) and Asian leopard cat (**Figure 2B**) lineages, we see common patterns in the evolutionary history of the two lineages. In the Asian leopard cat lineage, we see clear delineations between the domestic cat (*Felis catus*), rusty-spotted cat (*Prionailurus rubiginosus*), and flat-headed cat (*Prionailurus planiceps*), with overlapping signal between the fishing cat (*Prionailurus viverrinus*) and Asian leopard cat (*Prionailurus bengalensis*). The convoluted mixing of signal is likely explained by recurrent events of ancient hybridization given their broad overlap in geographic range Similar patterns of overlapping signal can also be seen within the Lynx lineage data set, specifically between the Iberian (*Lynx pardinus*) and Eurasian lynx (*Lynx lynx*), also best explained by rampant interspecific hybridization which has led to many local trees that do not support the species tree (Abascal et al. 2016; Li et al. 2019). These regions can be further explored in the context of phylogenetic variation in TreeViewer.

### *Shared features across* the *THEx Dashboards*

THEx offers a wide array of graph customization. A toolbar at the top of each dashboard provides several templates, color palettes, and graph attribute options that allow users to change how the graphs look and feel (**Figure 3**). Graphs can also be edited, enabling users to customize titles and axis labels directly on the graph within the browser without the need to change the raw data files. In addition, changes are saved when graphs are exported, producing publication-ready plots. As a simple demonstration, we show two of many possible themes and color combinations on graphs that compare recombination rate quantiles and topology frequencies between the autosomes and X of the Asian leopard cat lineage (**Figure 3**).

**Figure 3.**
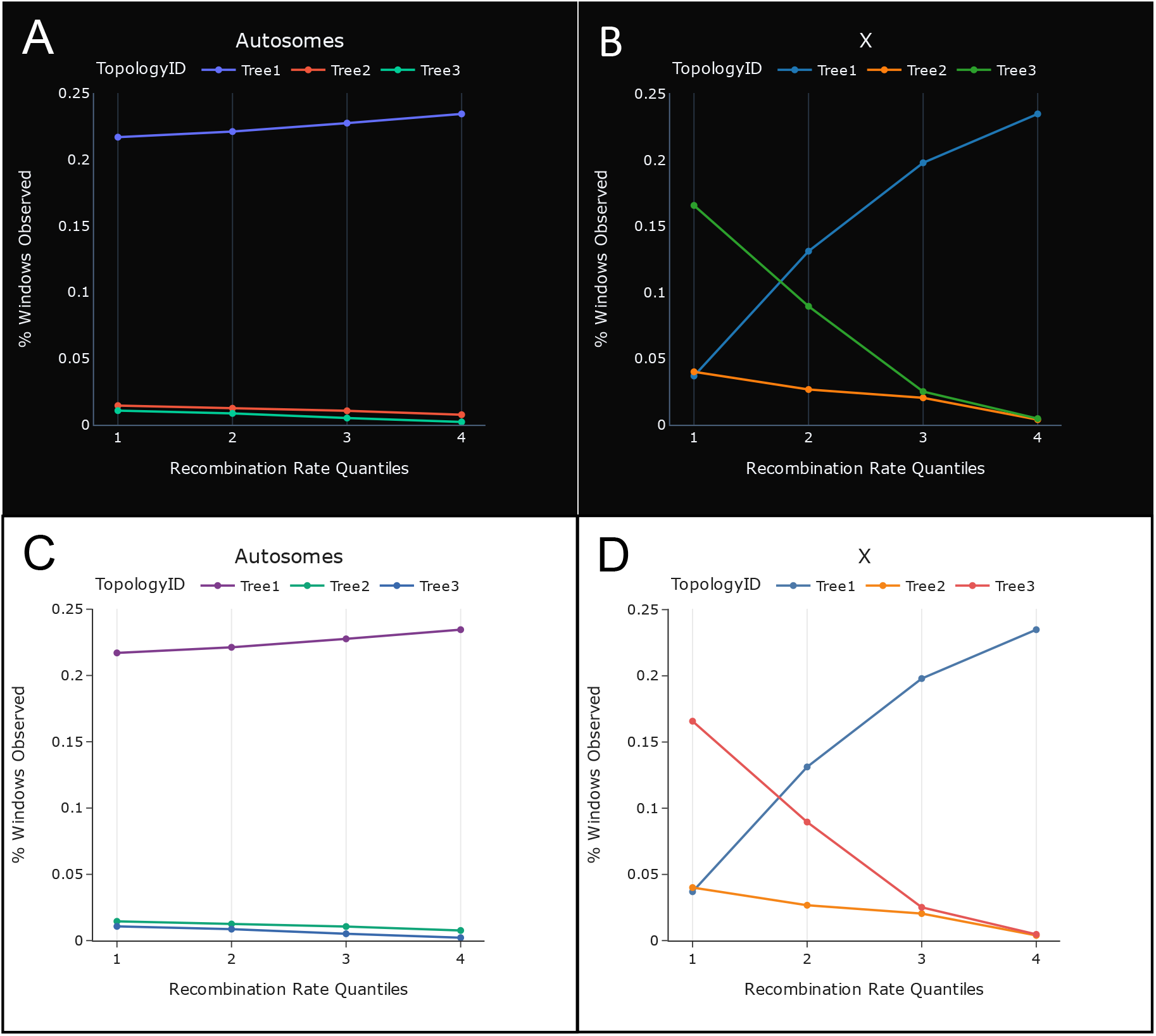
Comparison of the frequency of three tree topologies and recombination rate (cM/Mb) sorted low to high and partitioned into quantiles across the autosomes and X, respectfully, shows an increase in frequency of the true species tree (Tree 3) in low recombining regions of the X-chromosome. Paired graphs (A+B vs. C+D) demonstrate several graph template and color pallet combinations that can be interchanged on the fly within the graph customization menu of both Tree Viewer and Signal Tracer. These plots were produced using the Topology-Quantile tool under the Summary/Stats menu in Tree Viewer.

In addition to an extensive array of customization options, each graph is highly interactive, allowing users to hide or highlight specific data traces on the graph by selecting specific topologies (Tree Viewer) or species (Signal Tracer) labels within the legend. Clicking trace labels once in the legend removes the data traces from the graph and can be quickly added back in by clicking the same trace label again. Users can also single out individual traces by double-clicking legend items. This provides flexibility to simplify complex graphs by removing the need to produce multiple graphs for the same dataset.

Beyond graph customization and interactability, both Tree Viewer and Signal Tracer offer a “Current View” export option where users can download the underlying raw data from a specific genomic region. For example, a user zooms into a local region of a chromosome and finds an interesting divergent topology that aligns to a gene that is displayed from a loaded gene annotation (GFF/GTF) file. The user wishes to extract the information from the region being viewed from the Tree Viewer input file and gene annotation file. By clicking “File + Current View”, Tree Viewer will extract all information across all loaded data files for the given region and provide a download prompt for new input files with only the information residing within the region currently being viewed. This feature provides a rapid way to subset data and extract only the local information you want without the need to separately parse through large, whole genome data files.

## Results and Discussion

A growing number of studies (Edelman et al. 2019; Li et al. 2019; Small et al. 2020; Hennelly et al. 2021; Nelson et al. 2021) have demonstrated a strong correlation between recombination rate and the distribution of phylogenetic signal across the genome. In particular, Li et al. (2019) showed that across the X chromosome of all eight felid lineages, regions of low recombination were directly correlated with divergent regions of phylogenetic signal representing the true species tree. The major challenge presented in all of the analyses is the highly fragmented nature of the visualization of phylogenomic signal and direct comparison to corresponding data types, such as recombination. To demonstrate the power of the Tree Viewer dashboard within THEx, we re-analyzed the findings of Li et al. (2019) and extended some of their results by importing the raw data sets of the Asian leopard cat and Lynx lineages into a single genome browser. We utilize the Asian leopard cat data set to re-evaluate the distribution of signal for the true species tree across the X chromosome and identify novel signals of Canada lynx-bobcat hybridization within the lynx data set.

### Case Study 1: Phylogenetic discordance across the X-Chromosome in the Asian leopard cat lineage

We generated a Tree Viewer input file from the original Li et al. (2019) Asian leopard cat lineage raw data sets by taking the Newick trees labeled by a chromosome-window coordinate and organizing them into a tab-delimited file where the first three column headers were *Chromosome, Window*, and *NewickTree*. A Tree Viewer file requires one last column header, *TopologyID*, which can be generated by providing the original Newick tree files to PhyBin (Newton and Newton 2013) and passing the resulting directory to THExBuilder’s *“--phybin”* option, or providing the partially complete Tree Viewer input file to the “*--topobinner*’ option within THExBuilder to bin topologies in-house. Li et al. (2019) utilized PhyBin to cluster tree topologies across the genome based on a Robinson-Fould distance of 0 using the “*-bin*” parameter (Robinson and Foulds 1981). However, because PhyBin is not hosted on Conda, we created an alternate approach, *Topobinner*, that seamlessly integrates into Tree Viewer’s THExBuilder pipeline. We ran both PhyBin and Topobinner and verified that the “-- *topobinner*” option replicates the results from Li et al. (2019). We then added two additional data types, recombination rate and divergence time, by adding the values to the right of the first four required columns according to its corresponding chromosome-window coordinate. We initially investigated the phylogenetic signal from a whole-genome point of view as a rug plot, bar plot, pie chart, and 1-dimensional histogram faceted by chromosome, using Tree Viewer. We also explored the data on a per-chromosome basis, allowing us to zoom down to small local regions.

By adding recombination rate information and divergence time estimates, we could visualize and compare the recombination rate landscape across the X chromosome directly with phylogenetic signal (**Figure 1**). Through this comparison, we validated the results of Li et al. (2019), which concluded that signal for the true species tree resides in low recombining regions of the genome. For example, a ~40Mb recombination coldspot is found on the X chromosomes where ~66% of the windows reflect the species tree (Tree 3), while only ~17% of the windows support the most frequent genome-wide topology that most likely reflects signatures of gene flow or ILS (**Figure 1C**). Two smaller, multi-megabase regions of the X chromosome are similarly associated with low recombination and show the same trend of a large increase in the frequency of the true species tree (**Figure 1C-D**). Furthermore, when evaluating the divergence time estimates within these low recombining regions, older divergence times are found in the windows supporting the true species tree, again supporting the conclusions made by Li et al. (2019).

### Case Study 2: Signatures of Canada Lynx-Bobcat hybridization are correlated with decreased divergence time estimates

The Lynx lineage consists of four species: bobcat, Eurasian lynx, Iberian lynx, and Canada lynx (**Figure 4A**). The bobcat diverged from the common ancestor of the three lynx species approximately 3 million years ago and has a current geographic range that stretches from Central America to the southern border of Canada (Li et al. 2019). Previous studies have shown that the geographic range of the bobcat has been expanding northward into southern Canada, increasingly overlapping the geographic range of the Canada lynx. Several studies have also demonstrated hybridization between the Canada lynx and bobcat along the US-Canada border (Schwartz et al. 2004; Homyack et al. 2008; Koen et al. 2014). Throughout the past several million years climate change during interglacial periods likely has contributed to the contraction of the southern boundary of the Canada lynx range, facilitating the expansion of the bobcat into Canada (Koen et al. 2014). Although rare, naturally occurring Canada lynx-bobcat hybrids have been identified at the edge of the Canada lynx’s southern range in Maine, Minnesota, and New Brunswick (Homyack et al. 2008; Koen et al. 2014).

**Figure 4.**
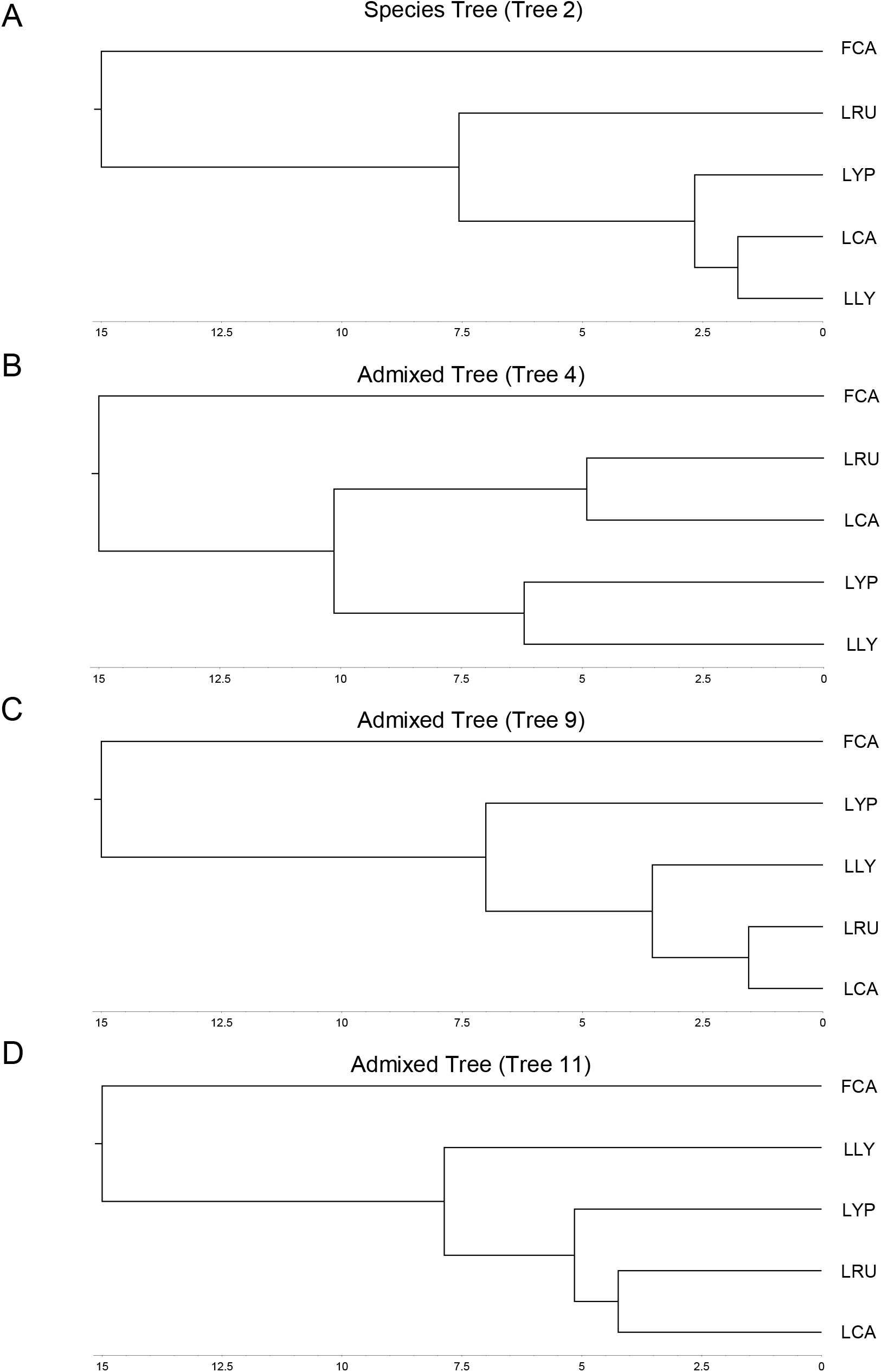
Time trees of the Lynx species tree (A) and Canada lynx-bobcat admixed topologies (B-D) with the Domestic Cat as the outgroup. Domestic cat (FCA), Canada lynx (LCA), Eurasian lynx (LLY), Iberian lynx (LYP), Bobcat (LRU).

To investigate potential signals of adaptive introgression that may have accompanied past northward range expansions of the bobcat during warmer, interglacial periods, we generated a Tree Viewer input file from the Lynx lineage data set from Li et al. (2019) and added divergence time estimates to the input file. We identified 42 windows (0.17%) with significantly lower (> 2 standard deviations) divergence time estimates and tree topologies that support post-speciation gene flow between the Canada lynx and bobcat (**Figure 4B-D, Figure 5**). Of the 42 outlier windows, three contained genes involved in adipogenesis and fat storage, all potentially important for adaptation to increasingly colder environments. The three windows are located on chromosome A3 and F2, windows 100.2Mb-100.3Mb and 30.4Mb, respectively (**Figures 6 & 7**). Spanning the two windows on chromosome A3, *RETSTAT* (Retinol Saturase) is a protein-coding gene known to cause increased adiposity in retinol saturase-knockout mice (Moise et al. 2010). On chromosome F2, several members of the *FABP* gene family are contained within the window supporting Canada lynx-bobcat hybridization and are all involved in fatty acid uptake, transport, and metabolism (Chmurzynska 2006).

**Figure 5.**
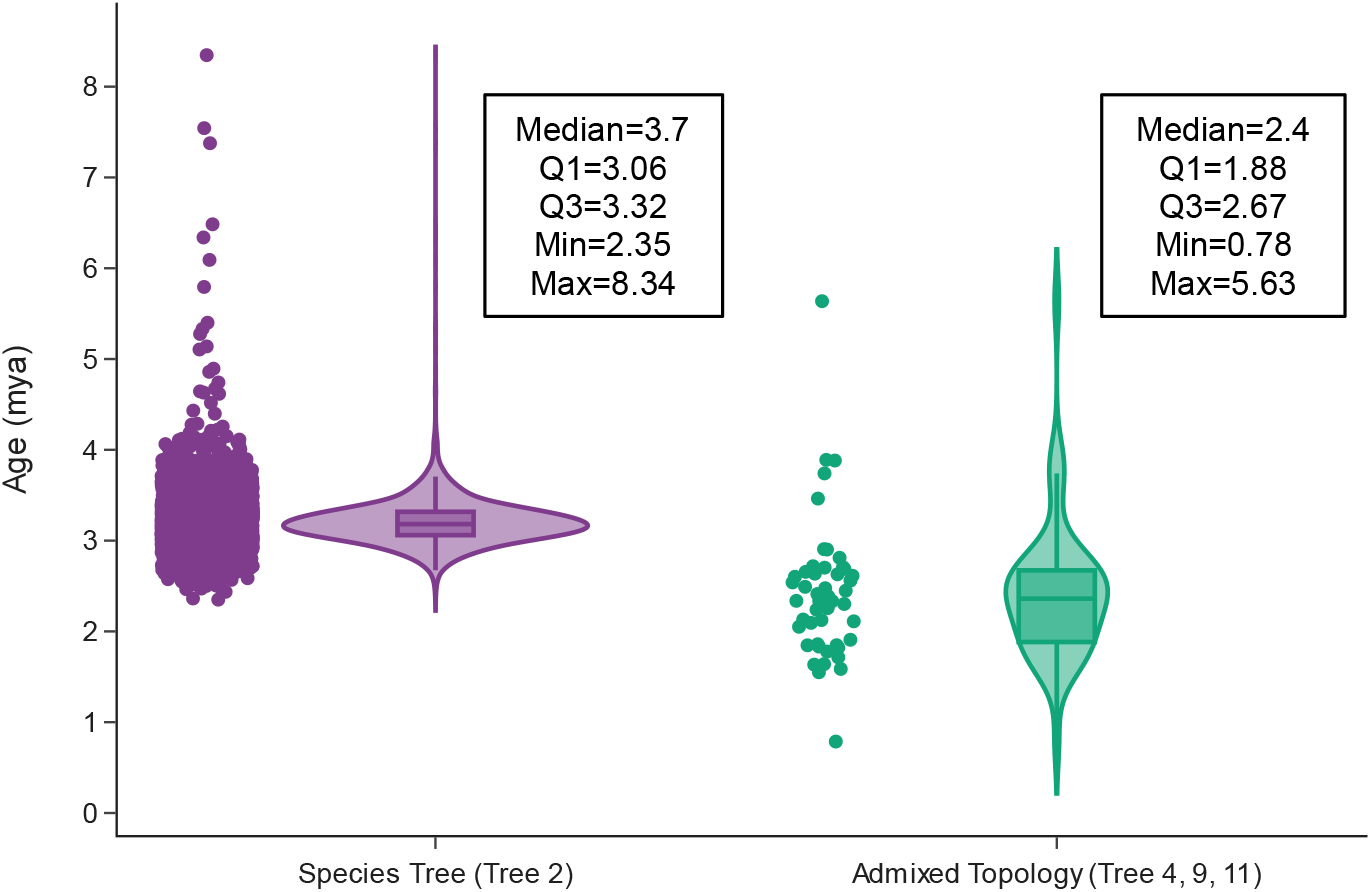
Divergence time estimates of bobcat in windows with topologies reflecting the species tree (Tree 2) and admixed trees (Tree 4, 9, 11). Divergence times from admixed topologies reflect significantly more dispersed and younger estimates, in and support of post-speciation gene flow between the Canada lynx and bobcat.

**Figure 6.**
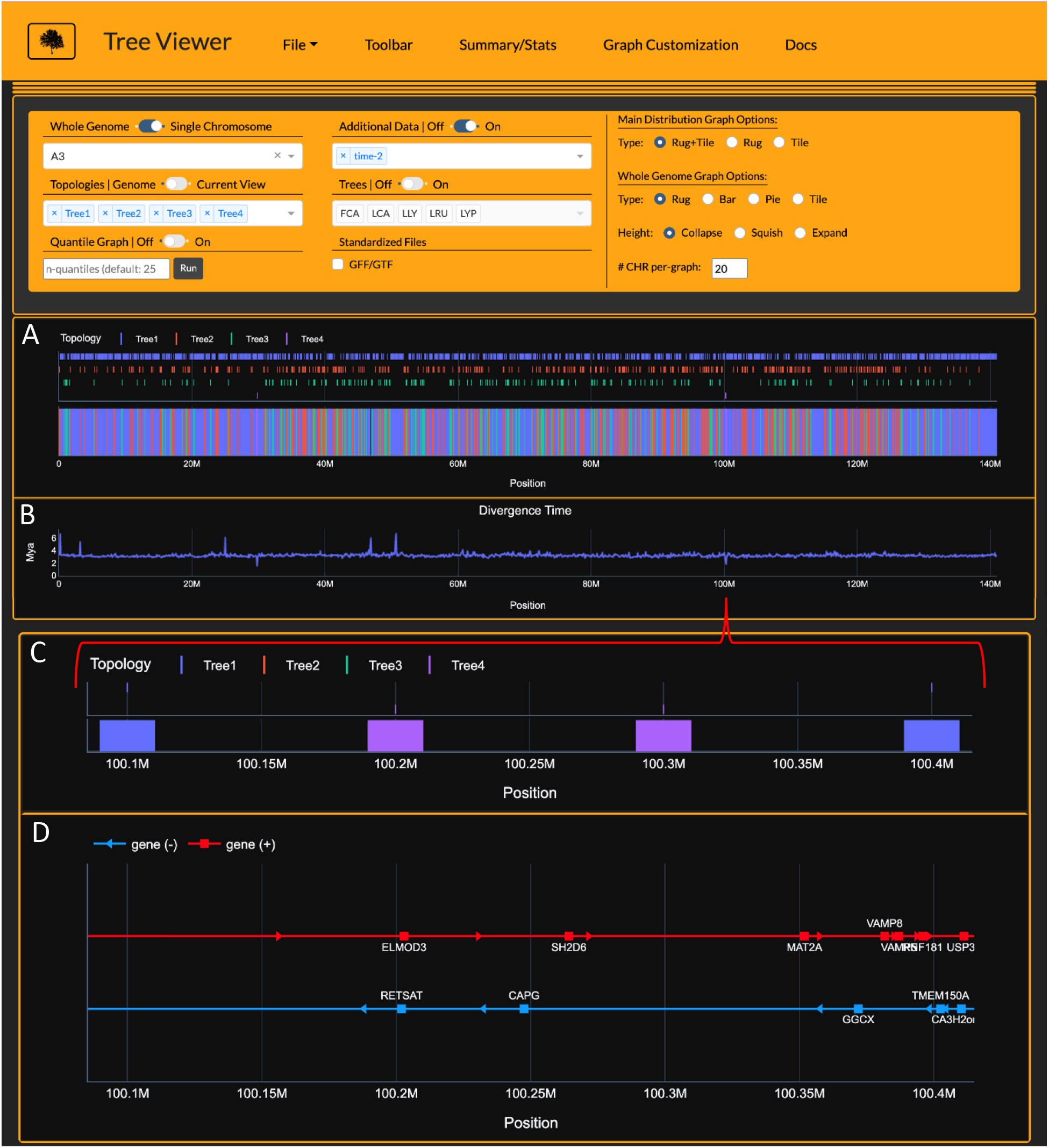
Tree Viewer interface depicting Lynx lineage phylogenetic signal (A), divergence time estimates (B), and zoomed-in view of Canada lynx-bobcat hybrid window (C) and underlying gene annotations (D) on chromosome A3. Divergence time estimates (time-2) are time estimates for branch 2 for each window’s respective tree topology. Importantly, time-2 estimates for Tree4 (Canada lynx-bobcat hybrid topology) indicate estimates for the bobcat, reflecting significantly younger divergence times at windows 29.8Mb, 100.2 Mb, and 100.3Mb (B). Gene annotations from windows 100.2 Mb and 100.3 Mb harbor *RETSTAT*, a gene known to cause increased adiposity in retinol saturase-knockout mice (Moise et al. 2010).

**Figure 7.**
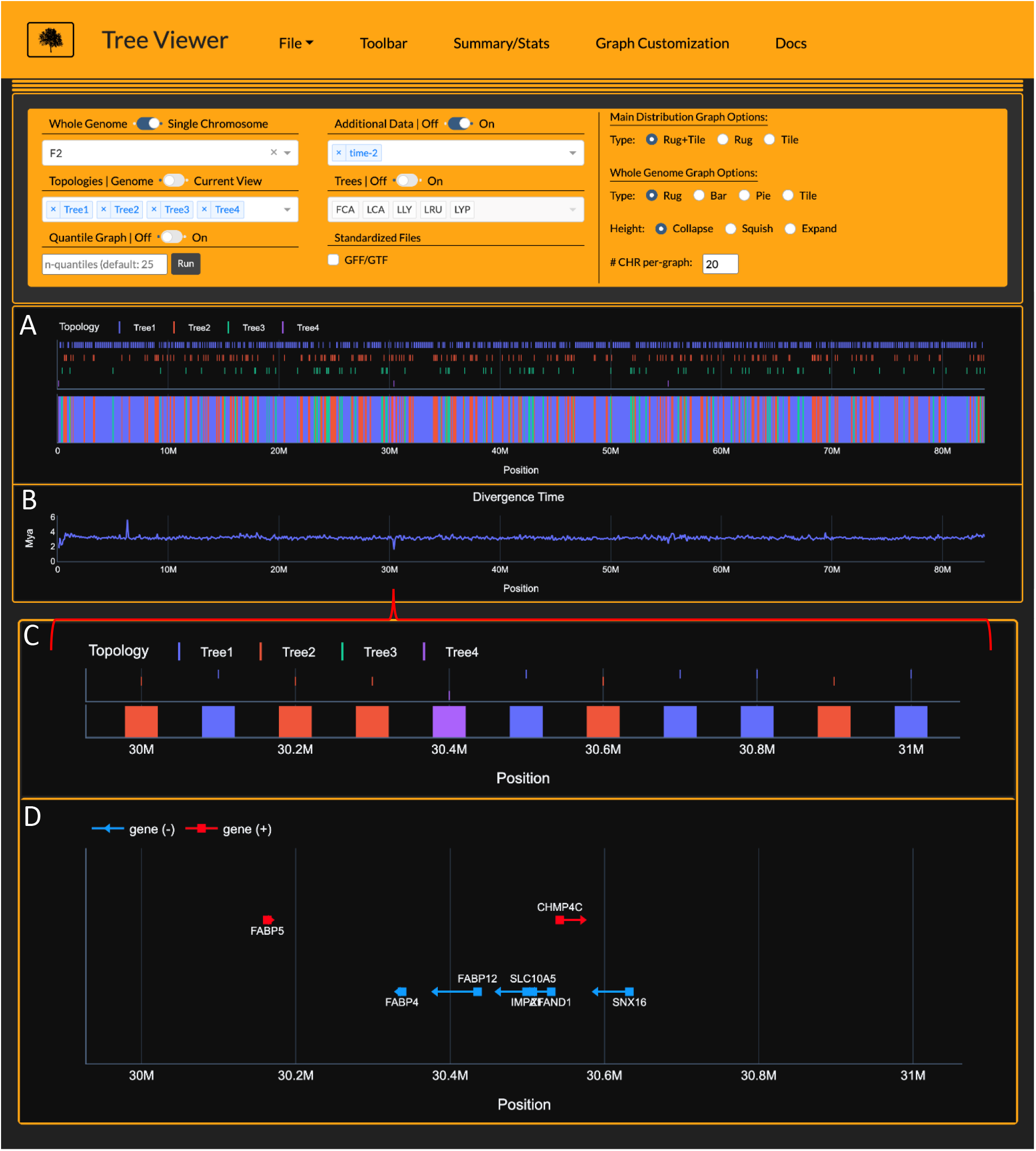
Tree Viewer interface depicting Lynx lineage phylogenetic signal (A), divergence time estimates (B), and zoomed-in view of Canada lynx-bobcat hybrid window (C) and underlying gene annotations (D) on chromosome F2. Divergence time estimates (time-2) are time estimates for branch 2 for each window’s respective tree topology. Importantly, time-2 estimates for Tree4 (Canada lynx-bobcat hybrid topology) indicate estimates for the bobcat, reflecting significantly younger divergence times at window 30.4 Mb (B). In addition, Gene annotations from window 30.4 Mb and adjacent regions (D) harbor the *FABP* gene family members involved in fatty acid take-up (Chmurzynska 2006).

Compared to the Canada Lynx, the bobcat is smaller in stature and typically occupies warmer environments, and lacks adaptations for deep snow. Bobcats also have poorer thermoregulatory capabilities in comparison to Canada lynx (Gustafson 1984; Mautz and Pekins 1989). We speculate that introgressed Canada lynx alleles of *RETSTAT* and the members of the *FABP* family may provide an adaptive advantage for northern bobcats to store larger quantities of fat and metabolize it more efficiently during the extremely cold months where prey availability is reduced. Further investigation of the underlying sequence properties and evolutionary rates at the population genetic level, as well as expression patterns of these candidate genes, are necessary to validate functional changes that would conclusively support the process of adaptive introgression. Nonetheless, this exercise exemplifies how the visualization of data using Tree Viewer can accelerate the genotype-phenotype discovery process and provide biological insights overlooked using previous approaches.

## Materials and Methods

THEx runs on Windows, macOS, and Linux operating systems and is hosted on Conda. Currently, THExBuilder only supports macOS and Linux as several pipeline dependencies lack support for Windows operating systems. Future development aims to resolve this limitation, enabling THExBuilder to run on all modern operating systems. Gevent, a coroutine-based Python networking library, provides the HTTP web service that communicates all data and user requests (interactions) to Plotly’s data analytic web application framework, Dash. Custom Python scripting adds functionality to THEx that is not natively provided in Dash and expands the functionality of existing components natively offered in Dash. All data used to exemplify the uses of THEx herein are available in the example directory on the THEx GitHub (https://github.com/harris-2374/THEx).

## Conclusion

THEx provides a novel and holistic approach to phylogenomic analyses that uniquely combines multiple data types to provide genomic context for the distribution of phylogenomic signal in a standalone application. THEx provides highly interactive graphing, making it easy to explore your data from a whole-genome, single chromosome, or local view. This approach facilitates a variety of analyses, including identification of the most probable species relationships, signatures of gene flow, or ILS, all in the context of genomic annotations. Designed to accommodate beginner and advanced computational biologists, THEx utilizes simple input file structures to allow users to generate them with any number of programs like Microsoft Excel or custom scripting. THExBuilder mitigates some of the more challenging aspects of input file creation and manipulation by providing a command-line suite of tools and pipelines. Together, THEx and THExBuilder provide a complete pipeline taking users from raw multiple-sequence alignments to phylogenomic analysis and integrated visualization, all within a single platform.

## Acknowledgments

We thank Jonas Lescroart, Kasuni Daundasekara, and Dr. Heath Blackmon for their contributions during the beta phase of development. Their input greatly improved the interactability and overall content within THEx.

